# Smoothened and ARL13B are critical in mouse for superior cerebellar peduncle targeting

**DOI:** 10.1101/2021.01.29.428892

**Authors:** Sarah K. Suciu, Alyssa B. Long, Tamara Caspary

## Abstract

Patients with the ciliopathy Joubert syndrome present with physical anomalies, intellectual disability, and a hindbrain malformation described as the “molar tooth sign” due to its appearance on an MRI. This radiological abnormality results from a combination of hypoplasia of the cerebellar vermis and inappropriate targeting of the white matter tracts of the superior cerebellar peduncles. ARL13B is a cilia-enriched regulatory GTPase established to regulate cell fate, cell proliferation and axon guidance through vertebrate Hedgehog signaling. In patients, mutations in *ARL13B* cause Joubert syndrome. In order to understand the etiology of the molar tooth sign, we used mouse models to investigate the role of ARL13B during cerebellar development. We found ARL13B regulates superior cerebellar peduncle targeting and these fiber tracts require Hedgehog signaling for proper guidance. However, in mouse the Joubert-causing R79Q mutation in ARL13B does not disrupt Hedgehog signaling nor does it impact tract targeting. We found a small cerebellar vermis in mice lacking ARL13B function but no cerebellar vermis hypoplasia in mice expressing the Joubert-causing R79Q mutation. Additionally, mice expressing a cilia-excluded variant of ARL13B that transduces Hedgehog normally, showed normal tract targeting and vermis width. Taken together, our data indicate that ARL13B is critical for control of cerebellar vermis width as well as superior cerebellar peduncle axon guidance, likely via Hedgehog signaling. Thus, our work highlights the complexity of ARL13B in molar tooth sign etiology.

**Summary statement:** Joubert syndrome is diagnosed by the hindbrain “molar tooth sign” malformation. Using mouse models, we show loss of the ciliary GTPase ARL13B, mutations in which lead to Joubert syndrome, result in two features of the molar tooth sign: hypoplasia of the cerebellar vermis and inappropriate targeting of the superior cerebellar peduncles. Furthermore, we demonstrate that loss of vertebrate Hedgehog signaling may be the underlying disrupted mechanism as we extend its role in axon guidance to the superior cerebellar peduncles.

## INTRODUCTION

Joubert Syndrome and Related Disorders (JSRD) are recessive congenital disorders with a variety of symptoms including developmental delay, intellectual disability, abnormal respiratory rhythms, ataxia, oculomotor apraxia, polydactyly, craniofacial defects, retinal dystrophy, nephronophthisis, and hepatic fibrosis (Parisi *et al*. 2007, Bachmann-Gagescu *et al*. 2020). While the exact prevalence of JSRD is not known, published statistics range from 1:80,000 to 1:100,000, but these may be underestimates as suggested by a recent study (Brancati *et al*. 2010, Nuovo *et al*. 2020). The characteristic neuroanatomical feature of JSRD is the molar tooth sign (MTS), which is caused by hypoplasia of the cerebellar vermis and thickened, elongated superior cerebellar peduncles (SCPs) that fail to decussate (Maria *et al*. 1997;Yachnis and Rorke 1999; Poretti *et al*. 2007). However, little is known about the mechanistic etiology of this hindbrain malformation. This is especially significant as several symptoms of JSRD arise from defects in the hindbrain: cerebellar dysfunction commonly causes ataxia and hypotonia, while some patients manifest life-threatening breathing problems (Parisi 2019).

To date, mutations in over 35 genes cause JSRD, and their associated proteins almost always localize to the primary cilium or the centrosome (Parisi 2019). Thus, JSRD is classified as a ciliopathy, a category of human disease stemming from ciliary dysfunction. One of the genes implicated in JSRD is *ARL13B*, which encodes a regulatory GTPase highly enriched in cilia (Cantagrel *et al*. 2008; Bachmann-Gagescu *et al*. 2015; Thomas *et al*. 2015; Shaheen *et al*. 2016; Rafiullah *et al*. 2017). As a GTPase, ARL13B is expected to have multiple effector proteins which interact with specific ARL13B residues. ARL13B can function as a guanine exchange factor (GEF) for ARL3, mutations in which also lead to JSRD (Gotthardt *et al*. 2015; Ivanova *et al*. 2017). JSRD-causing mutations in either ARL3 or ARL13B can disrupt their interaction or ARL13B’s GEF activity, consistent with the notion that specific ARL13B function is affected by JS-causing point mutations (Gotthardt *et al*. 2015; Ivanova *et al*. 2017; Alkanderi *et al*. 2018). Most JSRD-causing ARL13B mutations cluster within the protein’s GTPase domain, although two are located in the coiled coil domains in the C terminal half of the protein (Cantagrel *et al*. 2008; Bachmann-Gagescu *et al*. 2015; Thomas *et al*. 2015; Shaheen *et al*. 2016; Rafiullah *et al*. 2017). ARL13B complexes with the inositol phosphatase INPP5E, which is also implicated in causing JSRD (Bielas *et al*. 2009; Humbert *et al*. 2012). ARL13B is critical for targeting INPP5E to cilia and JSRD-causing ARL13B mutations disrupt INPP5E ciliary targeting (Humbert *et al*. 2012). INPP5E controls ciliary lipid composition through its phosphatase activity and most JSRD-causing mutations are within its phosphatase domain (Bielas *et al*. 2009; Chavez *et al*. 2015; Garcia-Gonzalo *et al*. 2015). Other proteins implicated in JSRD also affect ciliary targeting with many functioning at the transition zone, supporting the notion that abnormal ciliary traffic leading to defective signaling underlies JSRD (Arts *et al*. 2007; Delous *et al*. 2007; Garcia-Gonzalo *et al*. 2011; Hopp *et al*. 2011; Srour *et al*. 2012; Roberson *et al*. 2015).

The mechanistic connection between the cilia-related proteins implicated in JSRD and the MTS are elusive in part because distinct biological processes are at play. Abnormal proliferation may underlie the hypoplastic cerebellar vermis whereas defective axonal targeting is likely involved in the abnormal SCP tracts. One signaling pathway potentially linked to both processes is vertebrate Hedgehog (Hh) which relies on cilia (Huangfu *et al*. 2003). Sonic hedgehog (Shh) is a mitogenic cue that controls proliferation in the developing cerebellum so its misregulation could underlie the cerebellar hypoplasia (Dahmane and Ruiz i Altaba 1999; Wechsler-Reya and Scott 1999; Kenney and Rowitch 2000). While the SCP tracts that normally project from the deep cerebellar nuclei to the contralateral thalamus are guided by unknown signals, Shh is a known commissural axon guidance cue (Charron *et al*. 2003). JSRD patients can also display axon guidance defects in decussation of the pyramidal tracts (Yachnis and Rorke 1999; Poretti *et al*. 1997).

*ARL13B* and *INPP5E,* encoding ciliary proteins linked to JSRD, are known to regulate vertebrate Hh signaling. In mouse models, ARL13B loss disrupts cell fate specification in the neural tube, proliferation of the cerebellar granule precursor cells in the cerebellum and Shh-directed guidance of commissural axons in the spinal cord (Caspary *et al*. 2007; Bay *et al*. 2018; Ferent *et al*. 2019). These data support a model whereby disruption of Shh signaling by ARL13B mutation could provide a single mechanism underlying the MTS. This model is bolstered by the fact that additional phenotypes exhibited by JSRD patients, such as craniofacial defects or polydactyly, can arise from aberrant Hh signaling (Valente *et al*. 2008; Lan and Jiang 2009; Lipinski *et al*. 2010).

As attractive as a Hh-based model for JSRD may be, not all the data support that JSRD phenotypes result from misregulation of Hh signaling. Some features of JSRD, such as the renal and liver anomalies, are not clearly due to misregulation of Hh signaling (Doherty 2009; Breslow *et al*. 2018). Additionally, of over 35 genes implicated in JSRD, only 22 play some role in Hh pathway regulation (Davey *et al*. 2006; Reiter and Skarnes 2006; Caspary *et al*. 2007; Vierkotten *et al*. 2007; Huang *et al*. 2009; Weatherbee *et al*. 2009; Bimonte *et al*. 2011; Dowdle *et al*. 2011; Sang *et al*. 2011; Chih *et al*. 2011; Christopher *et al*. 2012; Thomas *et al*. 2012; Abdelhamed *et al*. 2013; Hynes *et al*. 2014; Wu *et al*. 2014; Chavez *et al*. 2015; Garcia-Gonzalo *et al*. 2015; Asadollahi *et al*. 2018; Frikstad *et al*. 2019; Munoz-Estrada and Ferland 2019). Some of these links are tenuous. For example, mouse *Arl3* mutants mislocalize the Hh transcription factor, GLI3, in their cilia yet do not exhibit any of the phenotypes normally displayed by mutants in the Hh pathway (Schrick *et al*. 2006, Lai *et al*. 2011; Schwarz *et al*. 2017).

Additional signaling pathways are linked to cilia including others known to be important in cell proliferation and axon guidance. Loss of either of the JSRD-linked genes *Ahi1* or *Cep290* in mouse leads to a small cerebellar vermis due to aberrant Wnt signaling (Lancaster *et al*. 2011; Ramsbottom *et al*. 2020). Possible JSRD-causing mutations in *Znf423* are associated with defects in Wnt, BMP and retinoic acid signaling (Hata *et al*. 2000, Huang *et al*. 2009, Casoni *et al*. 2020, Deshpande *et al*. 2020). Conditional *Arl13b* or *Inpp5e* deletion in the SCPs results in their disorganization and thickening through misregulation of ciliary PI3 kinase and AKT (Guo *et al*. 2019).

Ciliopathies are well established to be genetically complex. JSRD patients with different ARL13B mutations can display distinct phenotypes, such as obesity in an individual expressing ARL13B^Y86C^ and occipital encephalocele in a patient expressing ARL13B^R79Q^ (Cantagrel *et al*. 2008; Thomas *et al*. 2015). This is further exemplified in cases of related individuals carrying the same mutation but exhibiting different phenotypes and even diagnoses. For example, JSRD-causing mutations in TMEM67 (R208X) and TMEM216 (R73H) can also cause the more severe disease Meckel syndrome (Consugar *et al*. 2007; Otto *et al*. 2009; Valente *et al*. 2010). Understanding the genetic modifiers and environmental contribution underlying the phenotypic variation will be key to understanding disease etiology as will understanding when and how relevant pathways interact. In mouse models of the JSRD- and Meckel Syndrome-linked gene *Tmem67*, two phenotypic categories emerged: one with cerebellar malformations resembling JSRD and another with more severe CNS defects reminiscent of Meckel syndrome (Abdelhamed *et al*. 2013). The two categories correlated with whether cilia were retained, with the severe Meckel-like phenotype observed in animals lacking cilia. Furthermore, the two phenotypic groups impacted Hh and Wnt signaling differently, pointing to both pathways being critical (Abdelhamed *et al*. 2019). Importantly, it is not simply whether cilia are present, as another JSRD mouse model, *Talpid3*, lacks cilia yet displays a JSRD-like small cerebellar vermis (Bashford and Subramanian 2019). Thus, mouse models are incredibly informative yet point to the enormous complexity underlying the MTS.

Here we investigate the role of ARL13B in relation to Hh signaling in two major features of the MTS: targeting of the SCPs to the thalamus and hypoplasia of the cerebellar vermis. We explore these processes using a series of mouse alleles through which we first define the roles of Hh signaling and ARL13B in SCP projections. Subsequently, we untangle the role of ARL13B from within and outside of the cilium and investigate a JS-causing patient allele. Taken together, our data illuminate the roles of ARL13B in MTS etiology and the complexity in modeling aspects of the MTS in mouse.

## MATERIALS & METHODS

### Mouse lines

All mice were cared for in accordance with NIH guidelines and Emory University’s Institutional Animal Care and Use Committee (IACUC). Lines used were *Nex-Cre* (C3H-HeJ-*Neurod6^tm1(cre)Kan^*) [MGI:2668659], *Brn4-Cre* (C3H/HeJ-*Tg(Pou3f4- cre)32Cren*) [MGI:2158470], *Smo^flox^* (C3H/HeJ-*Smo^tm2AMC^*) [MGI:2176256], *Arl13b^flox^* (C3H/HeJ-*Arl13b^tm1Tc^*) [MGI:4948239], *Arl13b^V358A^* (C57BL/6J-*Arl13b^em1Tc^*) [MGI:6256969], and *Arl13b^R79Q^* (C57BL/6J-*Arl13b^em2Tc^*) [MGI:6279301]. Note that *Arl13b^Δ^* is the deletion allele resulting from germline deletion of the conditional *Arl13b^flox^* allele. Genotyping was as previously described (Heydemann *et al*. 2001; Goebbels *et al*. 2006; Nolan-Stevaux *et al*. 2009; Su *et al*. 2012; Gigante *et al*. 2020).

To generate the R79Q mutation in *Arl13b*, a CRISPR gRNA (ATTATTATGCTGAATCCTA*TGG*; PAM sequence is italicized) targeting exon 3 of the *Arl13b* locus along with a 180bp donor oligo (5’-CTCCCACTGTTGGCTTTTCTAAAA- TTGATCTGAGACAAGGAAAGTTCCAAGTTACCATCTTTGACTTAGGAGGTGGAAAA AGAATTC<u>A</u>GGG<u>C</u>AT<u>A</u>TGGAAGAATTATTATGCTGAATCCTATGGGGTAATATTTGTT GTGGATTCCAGTGATGAGGAGAGAATGGAAGAAACAAAGGAGA-3’; underlined bases are engineered) were designed to generate a G-to-A change creating the R79Q point mutation as well as A-to-C and T-to-A silent changes to create a *Nde*I restriction site that could be used for genotyping (Millipore Sigma). The gRNA (50 ng/ul), ssDNA oligo donor (50 ng/ul) and wild type *Cas9* mRNA (10 ng/ul) were injected into 1-cell C57BL/6J zygotes and subsequently transplanted at the 2-cell stage into C57BL/6J pseudopregnant females by the Emory Transgenic and Gene Targeting Core; all reagents were purchased from Millipore Sigma. Genomic DNA from toes was amplified via PCR using primers (5’-TCACTTGCAACAGAGCATCC-3’) and (5’- ACAGCTCTGCCCGTGTTTAC-3’) located upstream and downstream of the donor oligo breakpoints; products were sequenced with the forward (first) primer. A single founder animal heterozygous for both the R79Q mutation and NdeI restriction site was identified with no additional editing. Subsequent allele-specific genotyping of progeny was performed on ear punch or yolk sac using the following primers: Fwd-wt primer: 5’- GGAGGTGGAAAAAGAATaCg-3’; Fwd-mut primer: 5’- gctctatggctgGGAGGTGGAAAAAGAATTga-3’; Rev primer: 5’- AGTGCTAAGACACCCGAGGA-3’. PCR bands at 142bp (wild type) and/or 154bp (mutant) were produced, due to the addition of 12 non-templated bases to the 5’ end of the Fwd-mut primer (lowercase). Note the 3’ ends of the two forward primers differed in the base that codes for the R to Q change (final nucleotide of the primer) and includes a “wobble” base (lowercase) to provide allele-specific amplification after the first round of PCR (Gaudet *et al*. 2009). In order to breed away any potential off-target edits, the founder was backcrossed to C57BL/6J for three generations with at least two independent meiotic opportunities for recombination in each generation.

### Tract tracing injections and analysis

Tract tracing experiments were performed according to a protocol approved by Emory University’s Institutional Animal Care and Use Committee (IACUC). Male and female mice at postnatal day 90 or older were used for tract tracing experiments. At least 3 mice of each genotypic group were analyzed in experiments (exact N included in Figures 1-3, S1). Mice were anesthetized with inhaled isoflurane and maintained under anesthesia throughout the procedure. Animals were secured in the stereotax, and the scalp was opened with bregma and lambda aligned to flatskull position. Dorsal thalamus injections were targeted to the bregma (AP:-0.70, ML:+1.13, DV:-3.28, Angle:0°) and ventral thalamus injections were targeted using coordinates to the bregma (AP:-0.70, ML:-3.11, DV:-4.69, Angle:25°). Ventral injection targeting includes a 25° angle to avoid pulling dye through the dorsal thalamus upon needle removal. Then, a 5 ul Hamilton microsyringe was lowered to target and target was injected with lysine fixable dextran tetramethylrhodamine neuroanatomical tracer (fluoro-Ruby, 10,000 MW, ThermoFisher Scientific D1817). Animals received 0.05 - 0.5 ul injections of 10% dextran tetramethylrhodamine in sterile phosphate buffered saline (PBS, pH = 7.25) unilaterally at a rate of 0.1 ul/minute. The 0.5 ul injection volume initially used resulted in high background (non-specific) fluorescence. Later surgeries were conducted with 0.05 ul of dye, which resulted in lower background and retention of a strong signal in the DCN.

**Figure 1:**
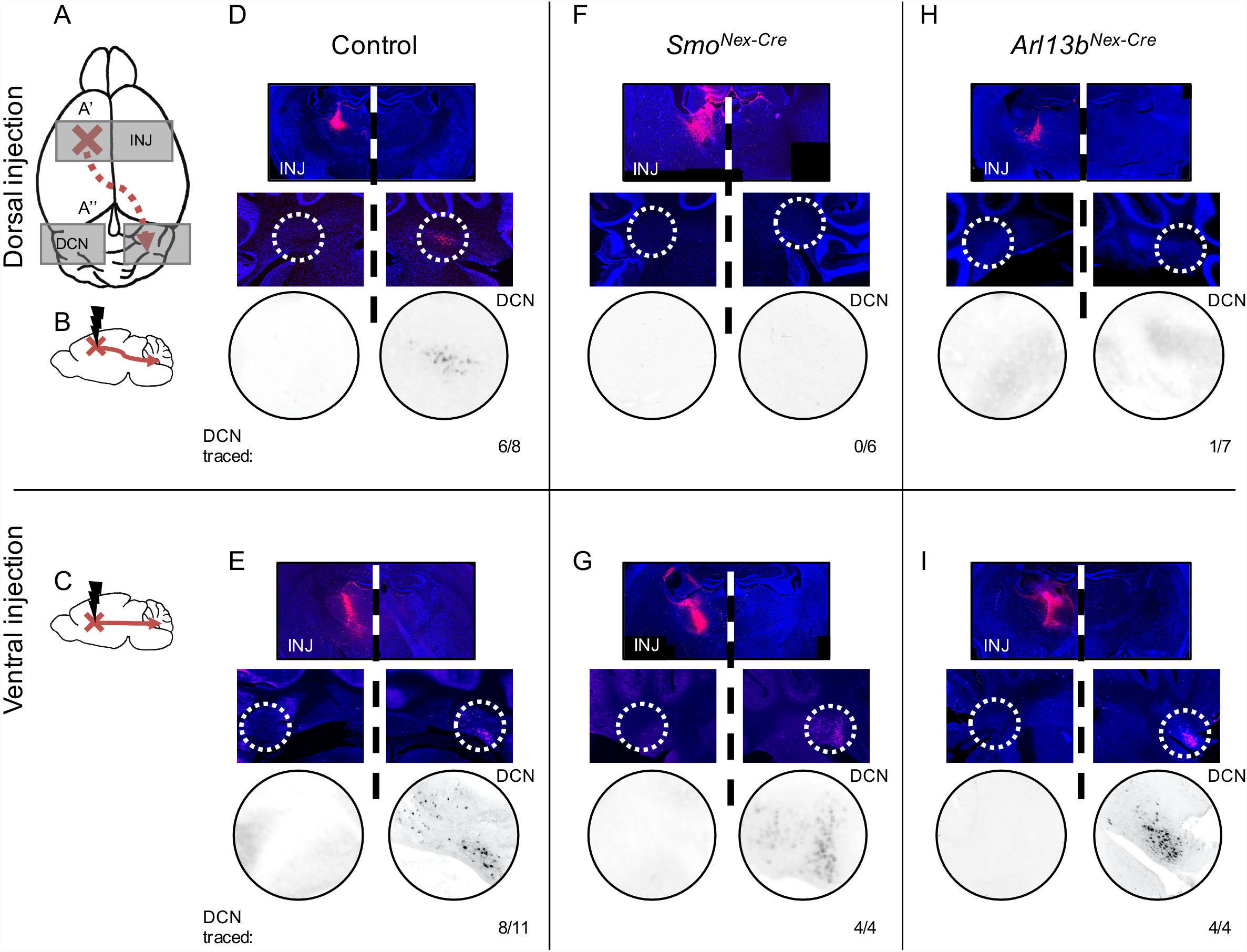
SCPs lacking *Arl13b* or *Smo* fail to project to the dorsal thalamus. (A-C) Schematics of injections and fluorescent tracer diffusion shown horizontally (A) or sagittally (B-C). (A) Red dashed arrow depicts dye path in a successful injection from injection site (red X) caudal through the brain and across the midline and into the contralateral cerebellar DCN (red arrowhead). Grey background boxes indicate area of subsequent images: the injection site (INJ) and cerebellum (DCN). (D-I) Representative images of dorsal (D, F, H) or ventral (E, G, I) thalamus injection site (top panel) and cerebellum (middle panel) with the DCN in hatched white circle and magnified (bottom panel) with recoloring to black and white to aid visualization. The retrograde fluorescent tracer is pink-red and sections are stained with DAPI (blue). Numbers indicate the number of positively stained DCN clusters (DCN traced) out of the total number of injected animals. Note that no tracing was observed on ipsilateral side to injection. (D, E) Fluorescent tracer injection in *Smo^fl/+^; Arl13b^fl/+^; Nex-Cre* control animals resulted in contralateral DCN staining in (D) 6/8 dorsal thalamus injections and (E) 8/11 ventral thalamus injections. (F, G) Fluorescent tracer injection in *Smo^Nex-Cre^* animals resulted in contralateral DCN staining in (F) 0/6 dorsal thalamus injections and (G) 4/4 ventral thalamus injections. (H, I) Fluorescent tracer injection in *Arl13b^Nex-Cre^* animals resulted in contralateral DCN staining in (H) 1/7 dorsal thalamus injections and (I) 4/4 ventral thalamus injections. Two-tailed Fisher’s exact test was performed; only the dorsal injection results were significantly different from the controls (F: p<0.01; H: p<0.05)

Seven days post-procedure, mice were perfused with 50mls of PBS followed by 30mls of 4% paraformaldehyde (PFA). Brains were kept in PFA overnight and subsequently placed in 30% sucrose in 0.1M phosphate buffer (pH 7.3) for cryoprotection for at least 48 hours. Brains were embedded in Tissue-Tek OCT compound (Sakura) for coronal cryostat sectioning. Sections were 60 microns thick and processed through 70% ethanol dehydration and 0.1% sudan black autofluorescence quencher, rehydrated in PBS and treated with DAPI to stain nuclei prior to fluorescence imaging. Images were taken on a Lionheart FX automated microscope (Biotek) or at 5x magnification on a Leica DM6000B microscope (Leica) using SimplePCI imaging software (Hamamatsu). The Leica images were subsequently stitched together to reveal the entirety of the brain section in Fiji (Schindelin *et al*. 2012) or Photoshop (Adobe).

Surgical injection sites were assessed to ensure dye was present at the desired injection site. If the injection was off-target or dye at the injection site was not seen, samples were removed from analysis. Cerebellar images from injections were evaluated for DCN staining with assessor blinded to genotype. The number of injections that resulted in fluorescent DCN for each injection site (dorsal and ventral thalamus) were compared between mutant and control genotypes using a two-sided Fisher’s exact test (PRISM 8.2.0).

### Phenotypic analysis of embryos

Timed matings of mice were performed to generate somite-matched embryos at embryonic day 10.5 (E10.5). Embryos were dissected in cold PBS and processed for immunofluorescence staining as previously described (Constable *et al*. 2020). Primary antibodies used were: mouse anti-Shh (5E1, 1:10), mouse anti-Pax6 (PAX6, 1:100) (Developmental Studies Hybridoma Bank), rabbit anti-Olig2 (AB9610, 1:500, Millipore Sigma), and mouse anti-Arl13b (N295B/66, 1:1500, NeuroMab). Multiple sections from three embryos of each genotype were examined.

### Analysis of anatomical measures

Weanling age (P20-P24) male and female mice were sacrificed and brains were harvested and fixed in 4% PFA overnight at 4⁰. Ten sex-matched pairs of mouse brains were collected for each genotype; control animals were either wild type or heterozygous for the *Arl13b* point mutation or floxed allele (so *Arl13b^R79Q/+^*, *Arl13b^V358A/+^* or *Arl13b^flox/+^*) or lacked *Brn4-Cre* (for *Arl13b^Brn4-Cre^*). Brains were imaged on a tilted stage to present a surface view of the cerebellum, with a standard ruler in frame to confirm scale, using a dissecting microscope (Leica MZFLIII). Measurements were made in FIJI (Schindelin *et al*. 2012) with the investigator blind to genotype. Whole cerebellar width was measured at the widest part of the cerebellum, coinciding with lobule CI or CII. Vermis width was calculated by measuring the widest part of lobule VII (Deshpande *et al*. 2020).

Hemisphere and vermis heights were measured at the longest rostro-caudal point of the hemisphere or at the midline of the vermis (see Figure 5). Body and brain weights were measured on a standard lab scale. For each sex- and age-matched pair, the ratio of mutant to control measures were calculated and graphed. A one-sample t-test was performed after transforming the ratio data to a log(2) scale to normalize data distribution (PRISM 8.2.0). Using the vermis width measures from our control brains (mean = 2.6mm, SD = 0.18), a power calculation for a one-sided t-test estimated 90% power to detect a 10% difference with 9-10 paired samples for a nominal alpha=0.05 (https://clincalc.com/stats/samplesize.aspx and https://www.stat.ubc.ca/~rollin/stats/ssize/n2.html).

### Data availability

Mouse lines are available upon request. Supplemental material available at figshare. Figure S1 contains diagrams illustrating the complete results of the injection and tract tracing experiments. Figure S2 contains a schematic of the CRISPR strategy.

The authors affirm that all data necessary for confirming the conclusions of the article are present within the article and figures.

## RESULTS

### SMO is required for normal SCP projection to the dorsal thalamus

In order to test whether proper projection of SCPs requires Hh signaling, we compared the SCP tracts in mice in which we deleted the gene encoding the obligate Hh transducer, *Smoothened* (*Smo*), to control animals. As *Smo* null embryos die during embryogenesis, we deleted *Smo* specifically in the projection neurons by generating *Nex-Cre;Smo^fl/fl^* mice, which we refer to as *Smo^Nex-Cre^* (Zhang *et al*. 2001; Caspary *et al*. 2002)*. Nex-Cre* initiates CRE recombinase expression at E11.5, as the precursor cells of the deep cerebellar nuclei (DCN) begin to migrate and become specified (Fink *et al*. 2006; Goebbels *et al*. 2006). In the mature cerebellum, the SCPs project rostrally from the DCN (illustrated in Figure 1A). After entering the midbrain, the SCPs cross the midline and again turn rostrally to project to two positions in the rostral thalamus: one tract takes a slight dorsal path and the other tract remains in the same plane; for simplicity here, we term these projection sites the dorsal and ventral thalamus, respectively (Bohne *et al*. 2019). To examine the SCP tracts, we used retrograde tract tracing in which we performed stereotaxic injections of a lipophilic dextran dye into either the dorsal (Figure 1B) or ventral (Figure 1C) thalamus and allowed the dye to diffuse through the axons to the associated neuron’s cell body (∼7 days); we then sacrificed the animal and examined the cerebellum for evidence of the lipophilic dye indicating tracing.

We found that both dorsal and ventral thalamus injections resulted in visible clusters of dye-stained cells in the contralateral DCN, and not the ipsilateral DCN, in control animals indicating the retrograde tract tracing reliably labelled the SCPs in our hands (Figures 1D-E, S1A-B, dorsal: 6/8; ventral: 8/11). In the *Smo^Nex-Cre^* mice, the results differed depending on whether we injected in the dorsal or ventral thalamus (Figures 1F-G, S1C-D). In the ventral thalamus injections, we detected dye-stained clusters of cells in the contralateral DCN but not the ipsilateral DCN, indicating normal SCP projection to the ventral thalamus (Figure 1G, 4/4 injections). This indicates that at least some SCPs cross the midline. In the dorsal thalamus injections, we could not detect dye-stained clusters of cells in either the contralateral or ipsilateral DCN (Figure 1F, 0/6 injections) suggesting that SCPs lacking SMO do not project to the dorsal thalamus. These data implicate SMO as critical for proper projection of the SCPs to the dorsal thalamus.

### ARL13B is required for normal SCP projection to the dorsal thalamus

Given that ARL13B regulates vertebrate Hh signaling in a variety of contexts, we next assessed ARL13B’s role in proper SCP projection. In order to delete *ARL13B* specifically in projection neurons, we generated *Nex-Cre;Arl13b^fl/fl^* mice, which we refer to as *Arl13b^Nex-Cre^.* We performed dorsal and ventral thalamus injections for retrograde tract tracing to examine the SCPs (Figures 1H-I, S1C-D). In the ventral thalamus injections of *Arl13b^Nex-Cre^* mice, we found dye-stained clusters of cells in the contralateral DCN consistent with normal SCP projections crossing the midline and projecting to the ventral thalamus (Figure 1I, 4/4 injections). In contrast, in the dorsal thalamus injections of *Arl13b^Nex-Cre^* mice, we generally did not detect dye-stained clusters of cells in either the contralateral or ipsilateral DCN, suggesting that the SCPs lacking ARL13B do not project to the dorsal thalamus (Figure 1H, 1/7 injections). These data link ARL13B function to normal SCP projection. Furthermore, they reveal a similar phenotype in *Smo^Nex-Cre^* and *Arl13b^Nex-Cre^* mice.

### ARL13B does not function from within cilia to mediate SCP guidance

*ARL13B* and the other 35 genes implicated in Joubert syndrome associate with the cilium or centrosome leading to the assumption that protein dysfunction from these locales underlies JSRD phenotypes (Parisi 2019). To directly ask whether ARL13B mediates SCP guidance to the dorsal thalamus from within cilia, we examined a mouse expressing a cilia-excluded variant of ARL13B, ARL13B^V358A^ (Figure 2) (Gigante *et al*. 2020). We previously demonstrated that ARL13B^V358A^ retains all known ARL13B biochemical activity, is undetectable in cilia yet transduces vertebrate Hh signaling normally (Mariani *et al*. 2016; Gigante *et al*. 2020). We found that either dorsal or ventral thalamus injections resulted in visible clusters of dye-stained cells in the contralateral DCN in control (Figures 2A-B, S1G-H, dorsal: 3/3; ventral: 4/4) and *Arl13b^V358A/V358A^* (Figure 2C-D, dorsal: 4/4; ventral: 3/3) animals. In the context of the previous result showing that *Arl13b^Nex-Cre^* mice display abnormal SCP projections to the dorsal thalamus, these data demonstrate that ARL13B does not function from within cilia to regulate SCP projections.

**Figure 2:**
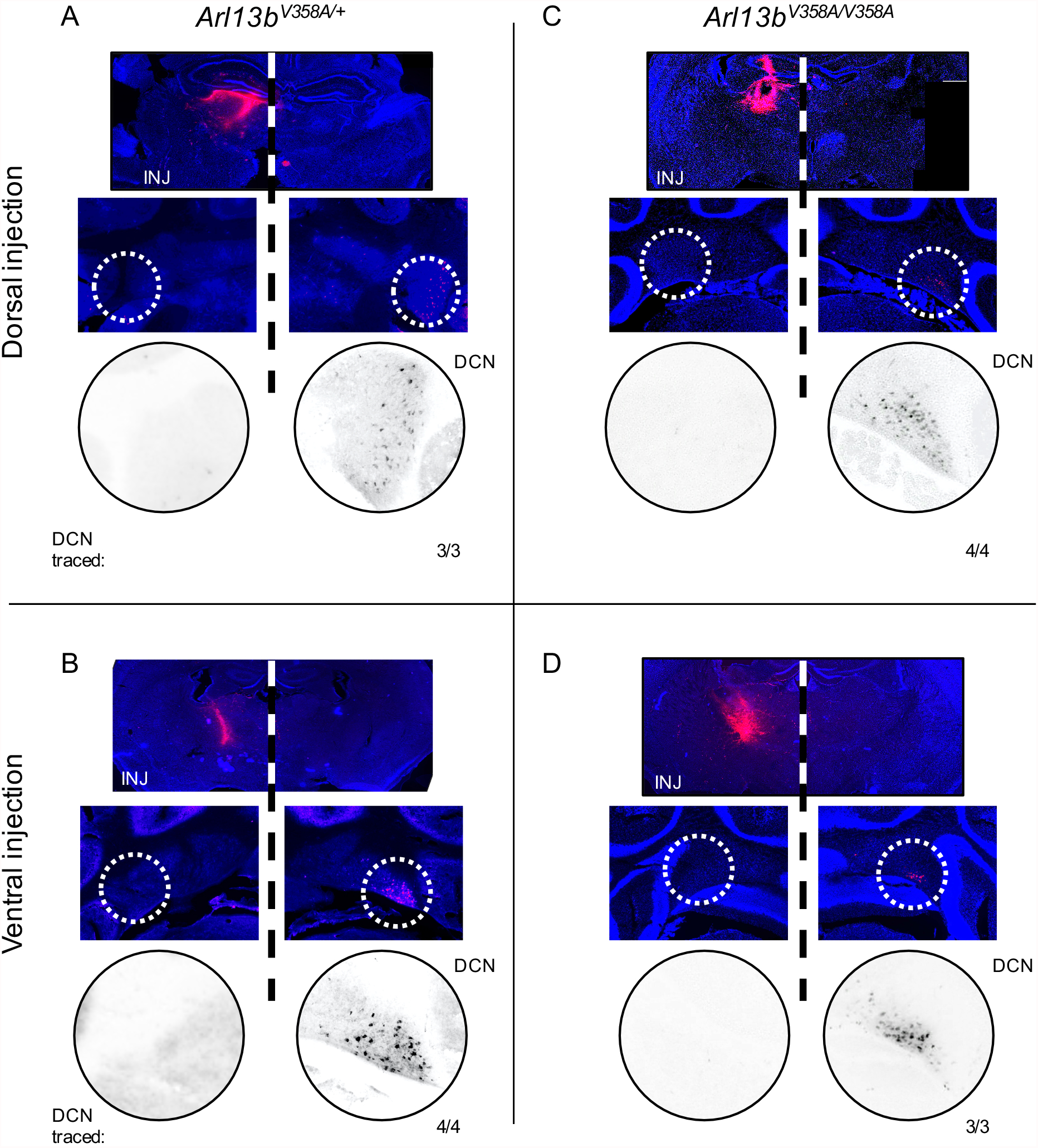
SCPs expressing cilia-excluded ARL13B^V358A^ project normally to both the dorsal and ventral thalamus. (A-D) Representative images of dorsal (A, C) or ventral (B, D) thalamus injection site (top panel) and cerebellum (middle panel) with the DCN in hatched white circle and magnified (bottom panel) with recoloring to black and white to aid visualization. Numbers indicate the number of positively stained DCN clusters out of the total number of injected animals. Note that no tracing was observed on the injection’s ipsilateral side. (A, B) Fluorescent tracer injection in *Arl13b^V358A/+^* control animals resulted in contralateral DCN staining in (A) 3/3 dorsal thalamus injections and (B) 4/4 ventral thalamus injections. (C-D) Fluorescent tracer injection in *Arl13b^V358A/V358A^* animals resulted in contralateral DCN staining in (C) 4/4 dorsal thalamus injections (Fisher’s exact test, two-tailed, ns, p>0.9) and (D) 3/3 ventral thalamus injections (ns, p>0.9).

### SCP projection is normal in mice expressing a Joubert-causing allele, Arl13b^R79Q^

In order to understand the relationship between ARL13B and MTS formation in Joubert syndrome, we generated a mouse expressing the JSRD-causing R79Q mutation (Figure 3A). We used CRISPR/Cas9 editing to change the conserved residue in the mouse genome (Figures 3B, S2). This amino acid change disrupts ARL13B’s GEF activity for ARL3 (Gotthardt *et al*. 2015; Ivanova *et al*. 2017). We found *Arl13b^R79Q/R79Q^* mice were viable and fertile. We bred the *Arl13b^R79Q^* allele to the null *Arl13b^Δ^* allele to make *Arl13b^R79Q^*^/^*^Δ^* animals, which we found survived to adulthood. As *Arl13b^Δ/Δ^* are embryonic lethal, this genetically demonstrates that *Arl13b^R79Q^* is a hypomorphic allele of *ARL13B* (Su *et al*. 2012).

**Figure 3:**
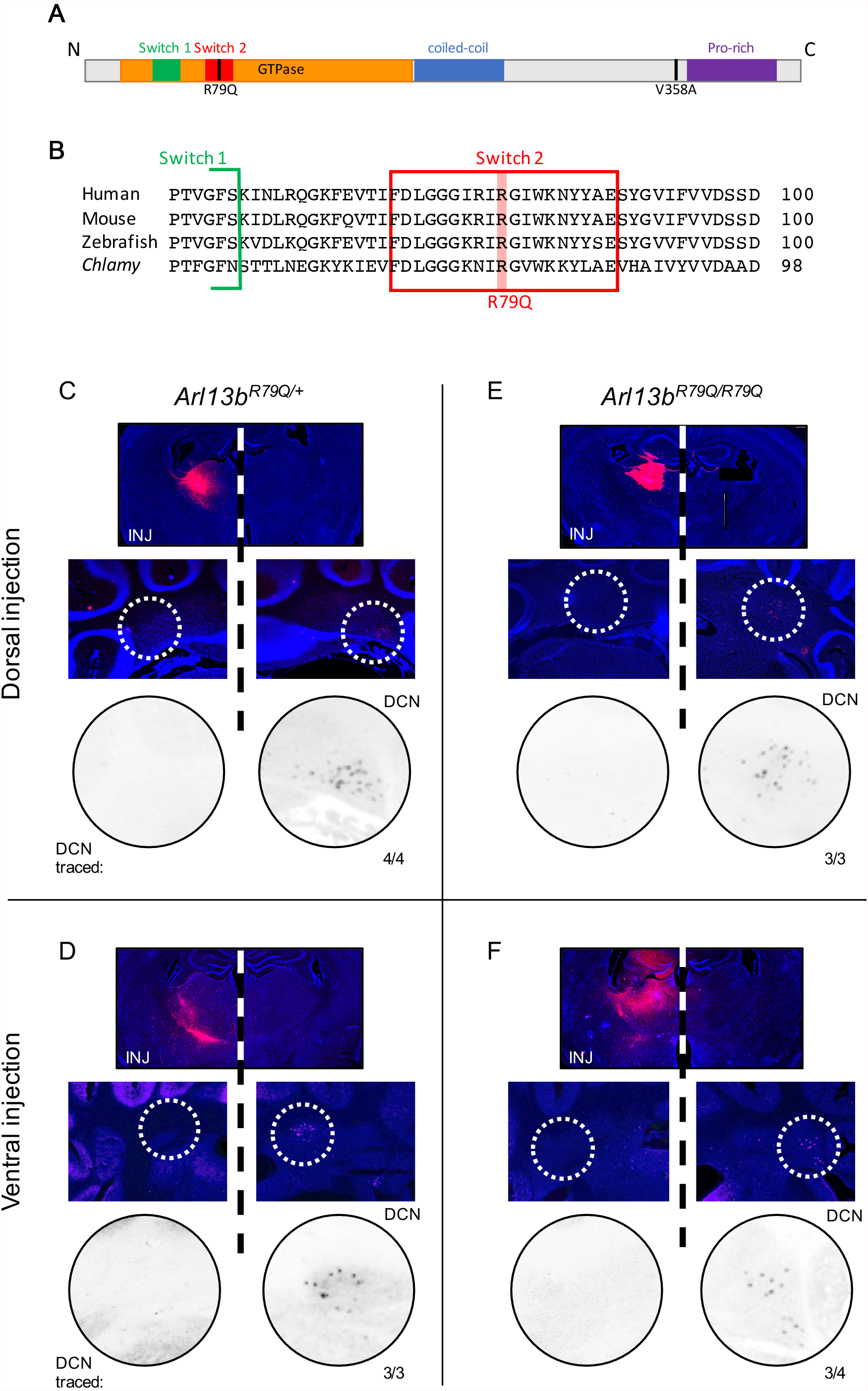
SCPs expressing JS allele *Arl13b^R79Q^* project normally to both the dorsal and ventral thalamus. (A) Schematic showing the protein domain structure of Arl13b. The R79Q mutation occurs in a highly conserved subregion of the GTPase domain, Switch 2 (red). (B) Alignment of protein sequence surrounding amino acid 79 of mouse Arl13b (red highlight) showing that arginine is conserved across multiple species. Sequences used in protein alignment: *Homo sapiens* NP_878899.1, *Mus musculus* NP_080853.3, *Danio rerio* NP_775379.1, *Chlamydomonas reinhardtii* XP_001691430.1. (C-F) Representative images of dorsal (C, E) or ventral (D, F) thalamus injection site (top panel) and cerebellum (middle panel) with the DCN in hatched white circle and magnified (bottom panel) with recoloring to black and white to aid visualization. Numbers indicate the number of positively stained DCN clusters out of the total number of injected animals. Note that no tracing was observed on the injection’s ipsilateral side. (C, D) Fluorescent tracer injection in *Arl13b^R79Q/+^* control animals resulted in contralateral DCN staining in (C) 4/4 dorsal thalamus injections and (D) 3/3 ventral thalamus injections. (E-F) Fluorescent tracer injection in *Arl13b^R79Q/R79Q^* animals resulted in contralateral DCN staining in (E) 3/3 dorsal thalamus injections (Fisher’s exact test, two-tailed, ns, p>0.9) and (F) 3/4 ventral thalamus injections (ns, p>0.9).

To assess the role of ARL13B^R79Q^ in SCP guidance, we performed dorsal and ventral thalamus dye injections in control and *Arl13b^R79Q/R79Q^* mice. We identified visible clusters of dye-stained cells in the contralateral DCN in control (Figures 3C-D, S1E-F, dorsal: 4/4; ventral: 3/3) and *Arl13b^R79Q/R79Q^* (Figure 3E-F, dorsal: 3/3; ventral: 3/4) animals. Thus, despite the constitutive expression of the JSRD-causing allele throughout development, we did not detect a SCP projection defect in the *Arl13b^R79Q/R79Q^* mouse model. In the context of the abnormal SCP projections to the dorsal thalamus that we identified in the *Smo^Nex-Cre^* and *Arl13b^Nex-Cre^* mice, this result indicates that the *Arl13b^R79Q^* allele does not disrupt SMO function or any ARL13B function that regulates SMO.

### Arl13b^R79Q/R79Q^ mice display normal Shh signal transduction in neural tube patterning

To further investigate the role of the *Arl13b^R79Q^* allele in Hh signaling, we examined embryonic neural patterning as it is exquisitely sensitive to alterations in Shh activity (Chiang *et al*. 1996; Briscoe and Ericson 1999). We generated E10.5 embryos and stained neural tube sections with antibodies against Shh, Olig2 and Pax6 (Figure 4). As expected in wild type embryos, we observed Shh expression in the ventral-most cells (the floorplate), Olig2 expression in lateral cells and Pax6 expression dorsally. We also saw the established abnormal cell fates in null *Arl13b^Δ/Δ^* embryos: loss of Shh staining in the floorplate (Figure 4D), dorsal and ventral expansion of Olig2 expression (Figure 4H), and a dorsal shift in Pax6 expression (Figure 4L) (Su *et al*. 2012). We found both *Arl13b^R79Q/R79Q^* (Figure 4B, F, J) and *Arl13b^R79Q/Δ^* (Figure 4C, G, K) embryos displayed neural patterning indistinguishable from wild type embryos, indicating the *Arl13b^R79Q^* allele is not dosage-sensitive and does not disrupt Shh signaling in determining neural cell fate. In addition, ARL13B^R79Q^ protein localized to cilia (Figure 4N), consistent with previous results (Li *et al*. 2010; Humbert *et al*. 2012; Li *et al*. 2016; Mariani *et al*. 2016).

**Figure 4:**
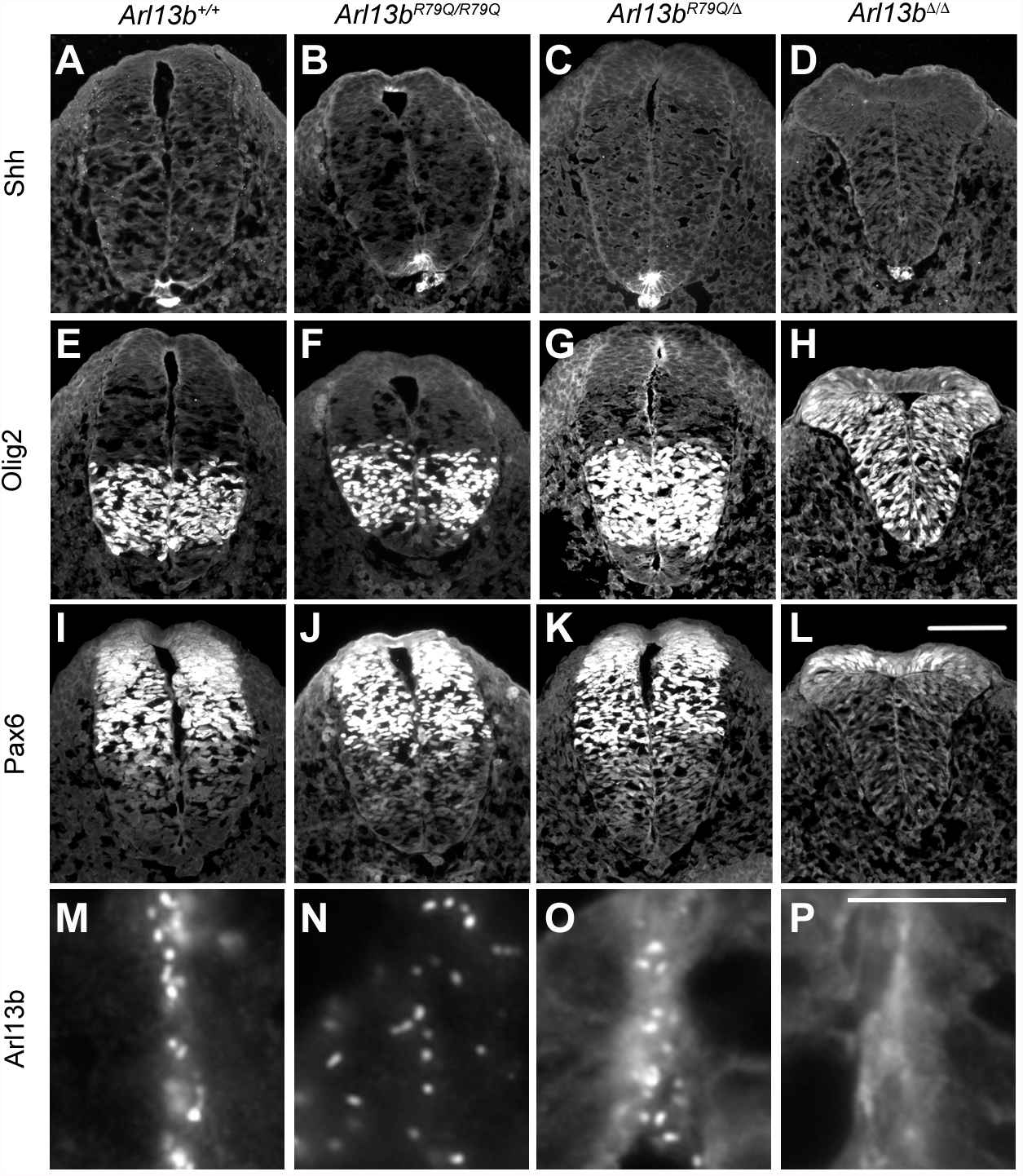
Mouse embryos expressing JS allele *Arl13b^R79Q^* display normal cell patterning in the neural tube. Shh, Olig2, Pax6 and Arl13b staining of E10.5 *Arl13b^+/+^* (n=3)*, Arl13b^R79Q/R79Q^* (n=3)*, Arl13b^R79Q/Δ^* (n=3), and *Arl13b^Δ/Δ^* (n=3) mouse neural tubes. Scale bar = 100 micrometers in A-L or 10 micrometers in M-P. (A-D) Shh is visible in the notochord and floorplate of (A) *Arl13b^+/+^,* (B) *Arl13b^R79Q/R79Q^* and (C) *Arl13b^R79Q/Δ^* neural tubes but absent from the floorplate of (D) *Arl13b^Δ/Δ^* embryos. (E-H) Olig2 stains the motor neuron precursor domain in (E) *Arl13b^+/+^*, (F) *Arl13b^R79Q/R79Q^* and (G) *Arl13b^R79Q/Δ^* embryos and stains an expanded domain in (H) *Arl13b^Δ/Δ^* embryos. (I-L) Pax6 expression is visible in the dorsal neural tube in (I) *Arl13b^+/+^*, (J) *Arl13b^R79Q/R79Q^* and (K) *Arl13b^R79Q/Δ^* neural tubes but shifted dorsally in (L) *Arl13b^Δ/Δ^* neural tubes. (M-P) Arl13b is localized to cilia visible in the ventral lumen of (M) *Arl13b^+/+^,* (N) *Arl13b^R79Q/R79Q^* and (O) *Arl13b^R79Q/Δ^* neural tubes but absent from (P) *Arl13b^Δ/Δ^* embryos.

### Arl13b^R79Q/R79Q^ mice display normal cerebellar width

The lack of a SCP projection phenotype in the *Arl13b^R79Q/R79Q^* mice surprised us since JS patients display the MTS. In addition to defects in the SCPs, the MTS is due to an underdeveloped cerebellar vermis, so we examined the width of the cerebellum and the cerebellar vermis (Figure 5) (Aguilar *et al*. 2012). To quantify cerebellar width, we performed analysis of surface-facing anatomical measurements validated to be sufficiently sensitive to detect small differences in vermis width (Deshpande *et al*. 2020). Briefly, we measured cerebellar width as well as cerebellar vermis width (widest part of lobule VII) of fixed whole mount dissected brains. For each sex- and age-matched pair, we calculated the ratio of the width measurements from mutant to control, which we compared to a hypothetical value of 1 (indicating no difference between groups). We detected no differences in the overall cerebellar width or the cerebellar vermis width between control and *Arl13b^R79Q/R79Q^* mice of either sex (Figure 5B). Furthermore, we found no difference in body or brain weight for *Arl13b^R79Q/R79Q^* mutants compared to controls (Figure 5B). Thus, unlike patients carrying *ARL13B^R79Q/R79Q^*, *Arl13b^R79Q/R79Q^* mice do not display a detectable growth deficit in the cerebellar vermis.

**Figure 5:**
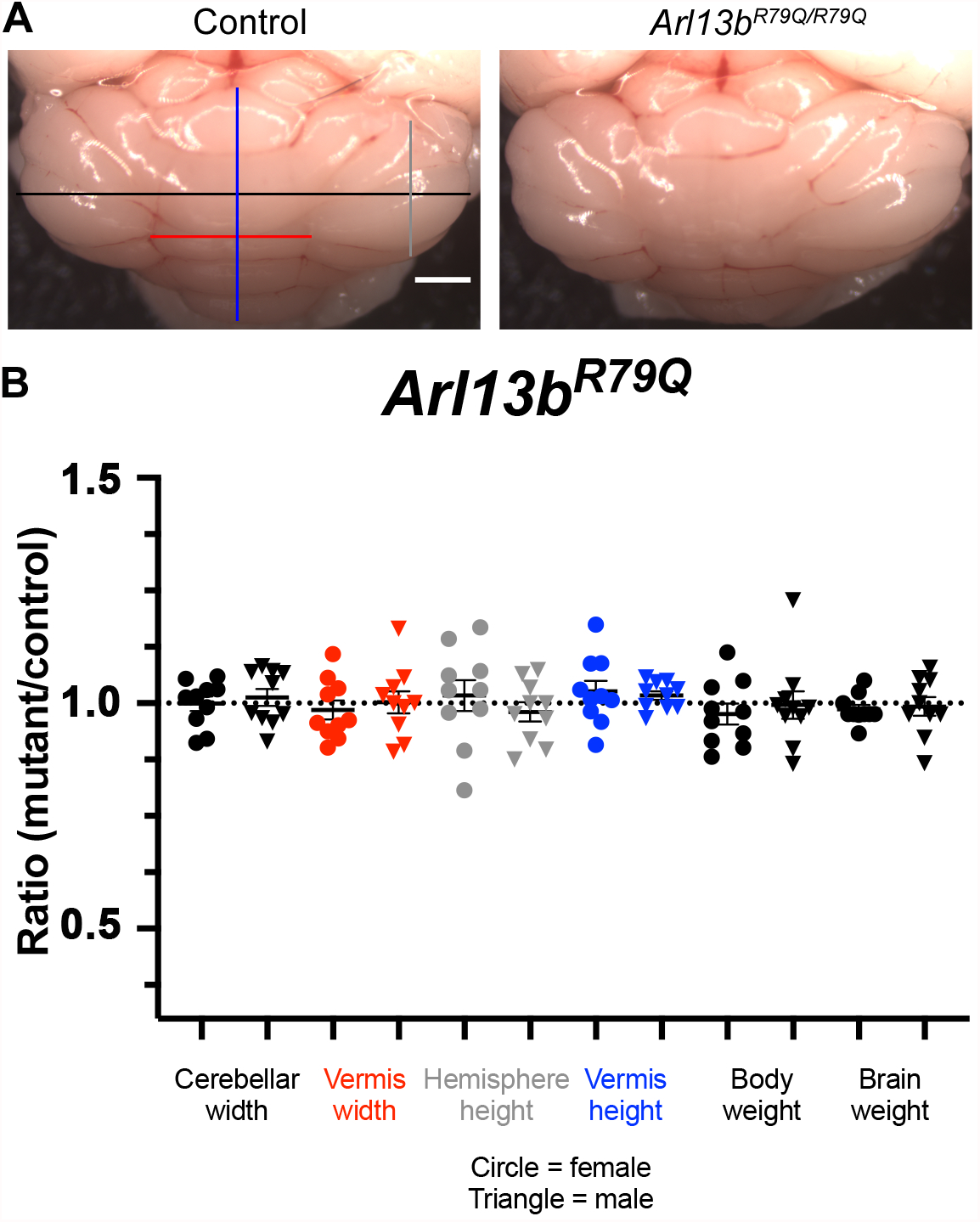
*Arl13b^R79Q^* mutation does not affect cerebellar width. (A) Representative surface-facing images of cerebella from control and *Arl13b^R79Q/R79Q^* mutant mice. Scalebar = 1mm, black line indicates where cerebellar width was measured, red line indicates where vermis width was measured, gray line indicates where cerebellar hemisphere height was measured, blue line indicates where vermis height was measured. (B) Ratios of *Arl13b^R79Q/R79Q^* mutant to control measurements of sex- and age-matched pairs of mice showed no significant difference. Cerebellar and vermis measurements were determined from surface views (symbol colors match the lines in A), body and dissected brain weights were determined using a standard lab scale. Each symbol represents a pair of sex- and age-matched animals: females are represented as circles and males are represented as triangles.

### Global cerebellar hypoplasia is observed in mice lacking Ar13b in all neurons

Cerebellar size is well established to be regulated, in part, via Shh signaling which controls proliferation of the cerebellar granule precursor cells (Kenney and Rowitch 2000; Chizhikov *et al*. 2007). In order to better understand the vermis hypoplasia phenotype seen in JBTS patients in relation to ARL13B, we wanted to investigate how ARL13B regulates cerebellar width. To do so, we crossed the *Brn4-Cre* allele into the conditional null *Arl13b^fl/fl^* background, called *Arl13b^Brn4-Cre^* (Figure 6). *Brn4- Cre* initiates expression at E8.5 throughout the neuroectoderm so the cerebellum develops in the absence of ARL13B (Heydemann *et al*. 2001; Hazen *et al*. 2012). We again calculated width ratios using surface-facing anatomical measurements and found the overall width of the cerebellum was 6% reduced in both females and males lacking ARL13B compared to control littermates at weaning (Figure 6B, p<0.1). More striking, in the cerebellar vermis we detected a 27% reduction in width in female and a 33% reduction in male mutants compared to controls (Figure 6B, p<0.0001). *Arl13b^Brn4-Cre^* mice develop hydrocephaly just after weaning which often leads to death. This is reflected in the body and brain weight ratios: while the mutant mice had slightly lower body weights, they had comparatively heavier brains (Figure 6B). From these data, we conclude that loss of *ARL13B* leads to a modest global cerebellar width deficit and a more pronounced cerebellar vermis width reduction.

**Figure 6:**
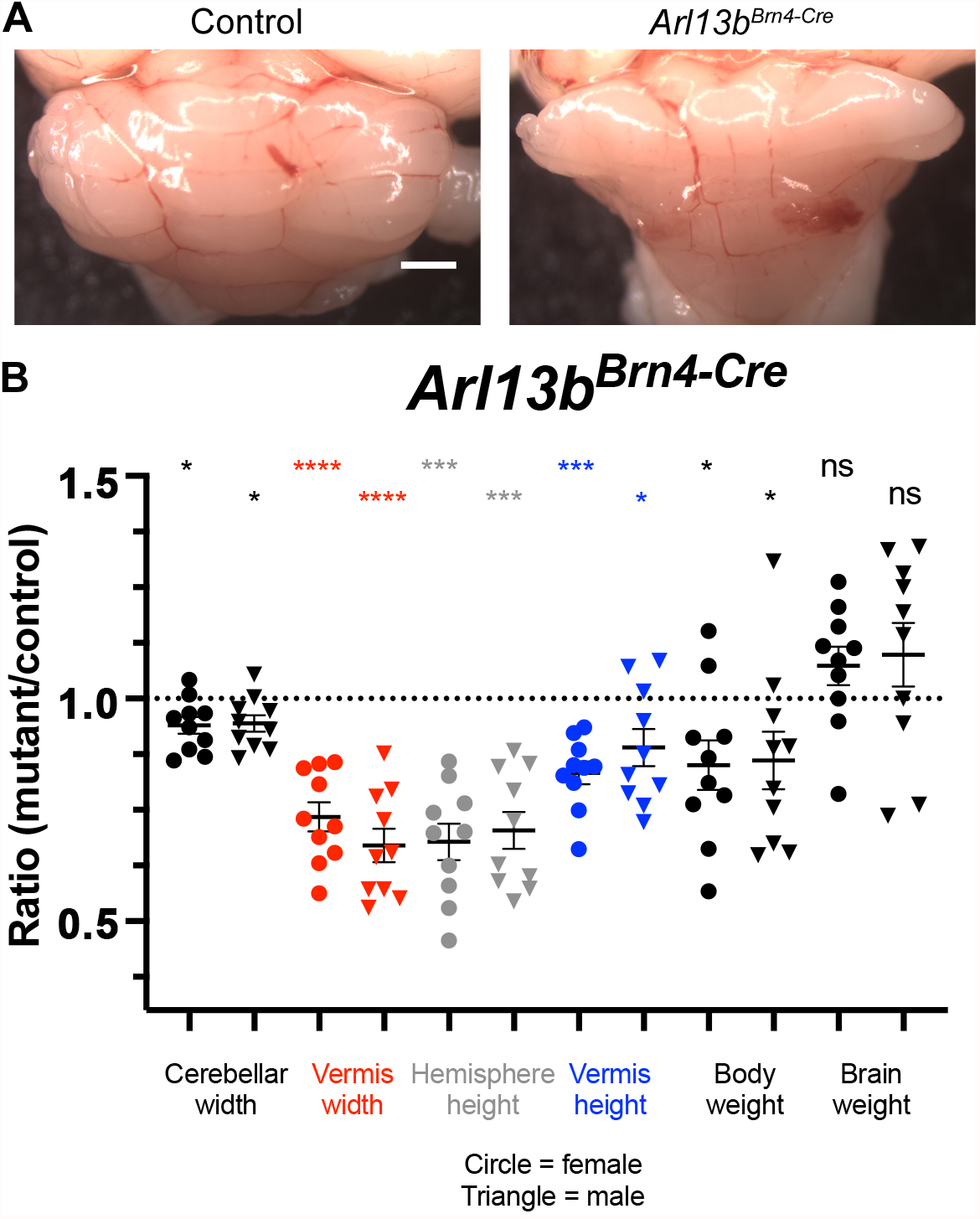
Pan-neuronal deletion of *Arl13b* results in a small cerebellum. (A) Representative images of cerebella from control and *Arl13b^Brn4-Cre^* mutant mice. Scalebar = 1mm. (B) The ratio of *Arl13b^Brn4-Cre^* mutant to control cerebellar width was reduced in both females and males (one-sample t-test: p<0.1, single asterisk). The ratio of *Arl13b^Brn4-Cre^* mutant to control vermis width was reduced in female and male age matched pairs (one-sample t-test: p<0.0001, four asterisks). While *Arl13b^Brn4-Cre^* mutants had moderately lower body weights than controls (one-sample t-test: p<0.1, single asterisk), their brains were slightly heavier, likely due to hydrocephaly. Each symbol represents a pair of sex- and age-matched animals: females are represented as circles and males are represented as triangles.

### Arl13b^V358A/V358A^ mice display normal cerebellar width

JS is classified as a ciliopathy due to the majority of causative genes encoding proteins that, like ARL13B, are associated with cilia. In order to better understand the role of ciliary ARL13B in cerebellar size, we examined cerebellar width in the mice expressing the cilia-excluded variant ARL13B^V358A^ (Figure 7) (Gigante *et al*. 2020). We detected no difference in the overall cerebellar width or that of the cerebellar vermis between control and *Arl13b^V358A/V358A^* mice (Figure 7B). Thus, ARL13B does not control cerebellar width from within cilia.

**Figure 7:**
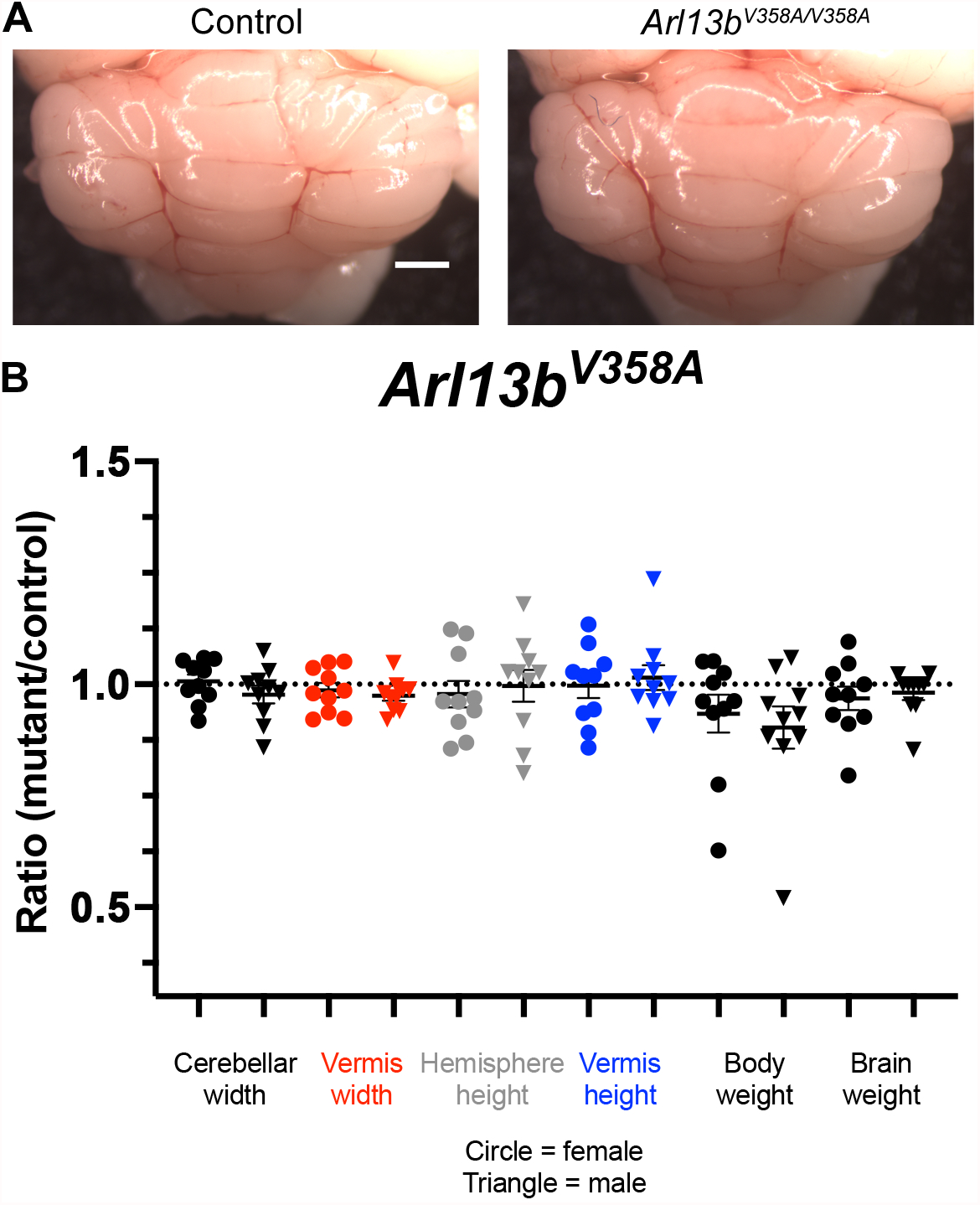
Mice expressing cilia-excluded *Arl13b^V358A^* have cerebella of normal width. (A) Representative images of cerebella from control and *Arl13b^V358A/V358A^* mutant mice. Scalebar = 1mm. (B) Ratios of *Arl13b^V358A/V358A^* mutant to control measurements of sex- and age-matched pairs of mice showed no significant differences (one-sample t-test). Each dot represents a pair of sex- and age-matched animals: females are represented as circles and males are represented as triangles.

## DISCUSSION

Here we demonstrate that complete loss of *Arl13b* function in mouse can account for two aspects of the MTS: aberrant SCP thalamic targeting and cerebellar vermis hypoplasia. We expand the role of Hh signaling as a critical guidance cue by showing it is required for proper SCP projection to the dorsal thalamus. Our finding that the SCP phenotype is similar in *Smo^Nex-Cre^* and *Arl13b^Nex-Cre^* mice is consistent with a model whereby ARL13B regulates SCP projections to the dorsal thalamus via a SMO-dependent mechanism. In line with previous work showing that ARL13B does not function from within cilia to regulate Shh-guided axon guidance, we found normal SCP thalamic targeting in mice expressing only a cilia-excluded ARL13B variant (Ferent *et al*. 2019). By mutating a conserved arginine to glutamine, we generated a mouse expressing a mutation linked to JSRD in humans and observed no change in vertebrate Hh signaling (Cantagrel *et al*. 2008). Additionally, we identified no defects in *Arl13b^R79Q/R79Q^* SCP projections. Whereas complete ARL13B deletion (*Arl13b^Brn4-Cre^*) in the cerebellum led to global hypoplasia, we show the cerebellum of *Arl13b^R79Q/R79Q^* mice is not significantly different in width compared to controls.

Overall, our data indicate ARL13B function is critical for both SCP targeting and controlling cerebellar vermis width. At one level, our data implicate Hh signaling in the etiology of the MTS since we show that SCP targeting requires SMO. However, at another level, our data indicate that Hh-independent pathways are at play as we don’t observe Hh-dependent neural tube patterning defects in the presence of the JSRD-causing *Arl13b^R79Q^* allele. In other mouse models of JSRD where the hypoplasia is specific to the cerebellar vermis, Wnt signaling is affected (Lancaster *et al*. 2011). Thus, the MTS could be due to disruption of different pathways in the SCPs and the vermis.

This would imply that the 35 JSRD implicated genes all affect the distinct pathways in a similar manner (Parisi 2019). Alternatively, the MTS may form due to alterations in any of a few pathways – and it is even possible that alterations in one pathway could impact other pathways – or the ability of cells to respond to those other pathways. Such a model is hinted at by previous work showing interplay between the Hh and Wnt pathways underlying the severity of hindbrain phenotypes (Hagemann and Scholpp 2012; Bashford and Subramanian 2019). While we haven’t detected any changes in Wnt response in the absence of ARL13B function, we may have not examined the relevant biological process or used a sensitive enough readout (Horner and Caspary 2011). Parallel reasoning would thus suggest that while *Arl13b^R79Q/R79Q^* mice clearly transduce Hh reasonably well, there may be subtle changes in Shh signaling or changes that influence Wnt signaling. Our data are consistent with the complexity exhibited by other JSRD mouse models examined to date (Delous *et al*. 2007; Garcia-Gonzalo *et al*. 2011; Roberson *et al*. 2015; Bashford and Subramanian 2019).

In patients, the SCP targeting deficit appears more severe than what we observed in the mice. Available methods for live-imaging and examination of post-mortem tissues suggest a range of SCP decussation defects, with some tracts appearing thickened on the ipsilateral side relative to their DCN (Yachnis and Rorke 1999; Poretti *et al*. 2007). Surprisingly, in the mutant mice we infer midline crossing of the SCPs. In the case of the *Smo^Nex-Cre^* conditional mice, it is formally possible that the SCPs do not rely on SMO for midline crossing but only for subsequent targeting to the dorsal thalamus. The fact that the *Arl13b^Nex-Cre^* conditional mice display a highly similar phenotype to the *Smo^Nex-Cre^* conditional mice makes this less likely, since ARL13B is directly implicated in JSRD and regulates SMO-dependent axon guidance in other contexts (Cantagrel *et al*. 2008; Ferent *et al*. 2019). It is also plausible that the protein turnover driven by *Nex-Cre* completed after midline crossing occurred. *Nex-Cre* expression initiates at E11.5 in the cells on the rhombic lip of the cerebellar anlage as they start to migrate and be specified before occupying the deep cerebellar nuclei (Fink *et al*. 2006; Goebbels *et al*. 2006). We expect deletion would occur in the precursors and therefore the neurons of the DCN would not express protein. Finally, it is possible that the mouse is not a valid system in which to model the SCP midline crossing defect.

This might explain why we saw no defects in the SCP targeting of the *Arl13b^R79Q/R79Q^* mice, as this is a constitutive mutation that requires no protein turnover, yet homozygous expression of ARL13B^R79Q^ in patients results in the molar tooth sign, which has been associated with failed decussation of white matter tracts (Quisling *et al*. 1999). Indeed, other mouse mutants such as *Cep290* and *Ahi1* which recapitulate the cerebellar vermis hypoplasia, also do not display midline crossing defects in the SCPs (Lancaster *et al*. 2011). Whether this is due to anatomical distinctions between the cerebellum in mouse and human or the genetic background on which these models were examined are open questions. Recent work highlights clear molecular and temporal differences between mouse and human cerebellar development (Haldipur *et al*. 2019; Behesti *et al*. 2021).

Examining SCP projections is labor intensive and it has not been done systematically among the JSRD mouse models (Bashford and Subramanian 2019; Guo *et al*. 2019). While previous work showed that *Arl13b^Nex-Cre^* and *Inpp5e^Nex-Cre^* mice exhibit SCP targeting deficits, here we pinpoint the *Arl13b^Nex-Cre^* defect as specific to the projection to the dorsal thalamus (Guo *et al*. 2019). The projection to the ventral thalamus remains intact, suggesting there is not a generalized deficit in axon outgrowth within the tract. The work on the *Arl13b^Nex-Cre^* and *Inpp5e^Nex-Cre^* SCP targeting deficits argue that PI3K/Akt signaling from within cilia led to the tract defects (Guo *et al*. 2019). However, we found that cilia-excluded ARL13B mediated SCP targeting normally.

These conflicting results could be explained by differences in the experimental details as the data supporting ciliary ARL13B function used a human ARL13B viral expression construct to rescue conditionally-deleted mice whereas we used genetic mutations engineered at the endogenous locus in this study. Alternatively, these data could indicate that ARL13B plays an important cellular role in the ciliary trafficking of key components needed for the PI3K/Akt pathways.

JSRD-causing mutations in ARL13B are generally restricted to the GTPase domain of the protein, although two residues outside that domain are implicated in disease (Cantagrel *et al*. 2008; Bachmann-Gagescu *et al*. 2015; Thomas *et al*. 2015; Shaheen *et al*. 2016; Rafiullah *et al*. 2017). Based on other ARL proteins, ARL13B likely assumes distinct conformations upon the binding either GDP or GTP, permitting different binding partners or altering affinities for binding partners (Pasqualato *et al*. 2002; Miertzschke *et al*. 2014). None of the tested JSRD-causing mutations (R79Q, Y86C or R200C) disrupt GTP binding or hydrolysis, however, all three mutations disrupt ARL13B function as an ARL3 GEF (Ivanova *et al*. 2017). The arginine at this position is located within the Switch 2 region of ARL13B and is conserved in humans, mice, zebrafish and *Chlamydomonas* (Figure 3B). We were surprised that we were unable to detect any defects in our mouse, as we expected to model some aspect of Joubert syndrome. Humans with homozygous R79Q mutation exhibit motor and ocular defects (among other symptoms), whereas *arl13b*-null zebrafish injected with human ARL13B- R79Q mRNA only partially rescue the ciliopathy phenotypes of curved body axis and cystic kidney (Cantagrel *et al*. 2008). These species-dependent differences emphasize the importance of analyzing mutations expressed from the endogenous promoter over the course of normal development. A recent study linked *arl13b* disruption during zebrafish development to reduced granule and Purkinje cells through a down-regulation of Wnt signaling (Zhu *et al*. 2020). Given that complete deletion of ARL13B impacts broader biological processes in the cerebellum than the R79Q mutation and that the null mutant misregulates Hh signaling whereas R79Q does not, we conclude that a subset of ARL13B function is disrupted in JSRD.

## Acknowledgements

We are grateful to L. Mariani for her initial work on this project, R.E. Van Sciver and E. Gigante for critical comments on the manuscript and J.G. Mulle for the statistical consults.

## Funding

This work was supported by funding from National Institutes of Health grants R01NS090029, R01GM110663 and R35GM122549 to T.C. and T32GM008490 and F31NS101806 to S.K.S. with additional support from the Emory University Integrated Cellular Imaging Microscopy Core of the Emory Neuroscience NINDS Core Facilities grant, P30NS055077. This study was supported in part by the Mouse Transgenic and Gene Targeting Core (TMF), which is subsidized by the Emory university School of Medicine and is one of the Emory Integrated Core Facilities. Additional support was provided by the National Center for Advancing Translational Sciences of the National Institutes of Health under Award Number UL1TR000454. The content is solely the responsibility of the authors and does not necessarily reflect the official views of the National Institutes of Health

## Conflicts of Interest

The authors have no competing interests to declare.

## Author contributions statement

Author Contributions: Conceptualization T.C.; Methodology S.K.S. and A.B.L.; Validation S.K.S. and A.B.L; Formal Analysis S.K.S. and A.B.L.; Investigation S.K.S. and A.B.L.; Writing – Original Draft S.K.S. and T.C.; Writing – Review & Editing S.K.S., A.B.L. and T.C.; Visualization S.K.S. and A.B.L.; Supervision T.C.; Project Administration T.C.; Funding Acquisition T.C.

## Notes

### Competing Interest Statement

The authors have declared no competing interest.

### Summary of Updates

Joubert syndrome is diagnosed by the hindbrain molar tooth sign malformation. Using mouse models, we show loss of the ciliary GTPase ARL13B, mutations in which lead to Joubert syndrome, result in two features of the molar tooth sign: hypoplasia of the cerebellar vermis and inappropriate targeting of the superior cerebellar peduncles. Furthermore, we demonstrate that loss of vertebrate Hedgehog signaling may be the underlying disrupted mechanism as we extend its role in axon guidance to the superior cerebellar peduncles.

