## Supplementgal figure 1 for "Smoothened and ARL13B are critical in mouse for superior cerebellar peduncle targeting"

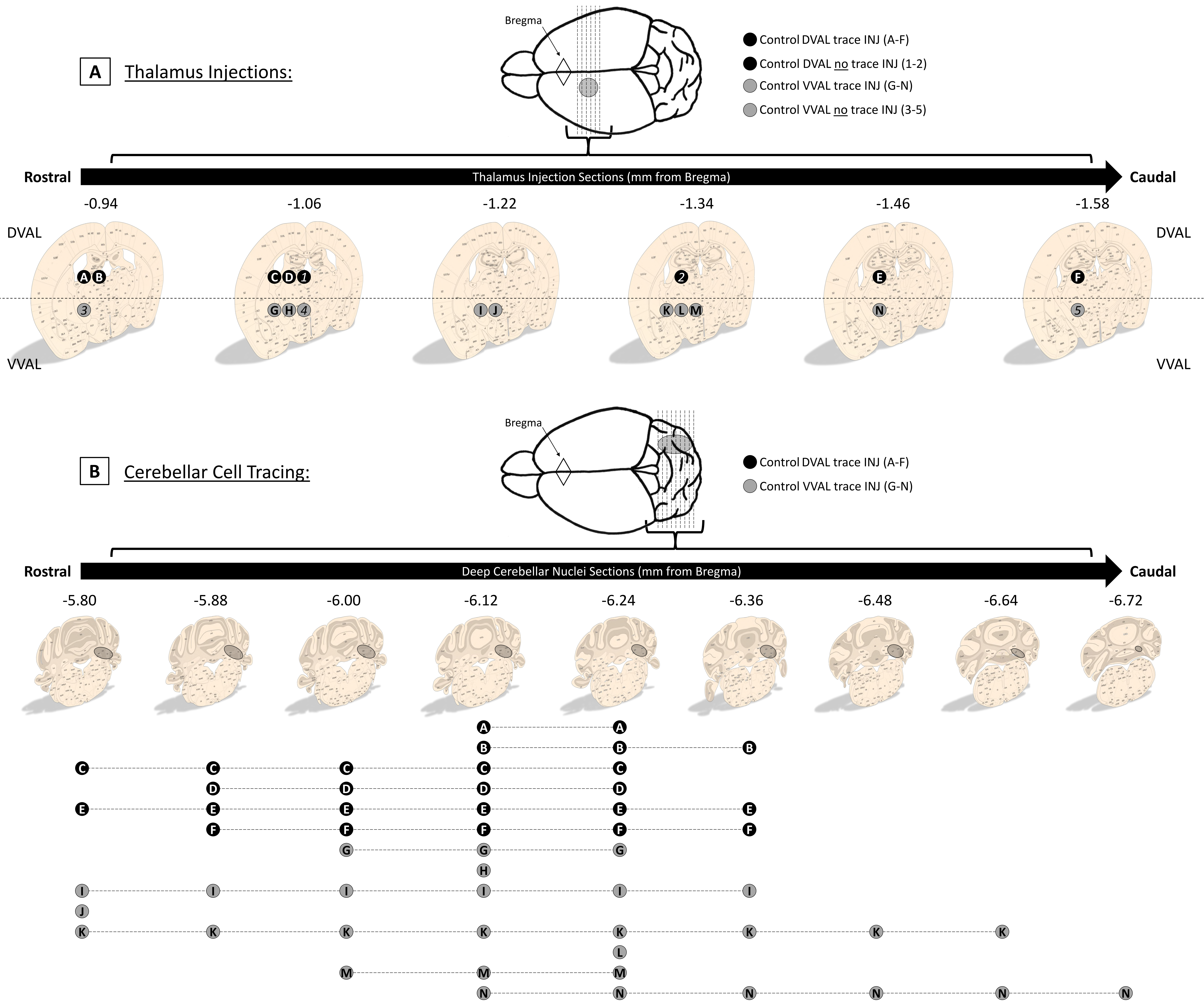
A
Thalamus Injections:
Bregma
Control DVAL trace INJ (A-F)
Control DVAL no trace INJ (1-2)
Control VVAL trace INJ (G-N)
Control VVAL no trace INJ (3-5)
Rostral
Thalamus Injection Sections (mm from Bregma)
Caudal
-0.94
-1.06
-1.22
-1.34
-1.46
-1.58
DVAL
VVAL
A B
C D 1
2
E
F
3
G H 4
I J
K L M
N
5
B
Cerebellar Cell Tracing:
Bregma
Control DVAL trace INJ (A-F)
Control VVAL trace INJ (G-N)
Rostral
Deep Cerebellar Nuclei Sections (mm from Bregma)
Caudal
-5.80
-5.88
-6.00
-6.12
-6.24
-6.36
-6.48
-6.64
-6.72
A A
B B B
C C C C C
D D D D
E E E E E E
F F F F F
G G G
H
I I I I I I
J
K K K K K K K K
L
M M M
N N N N N N

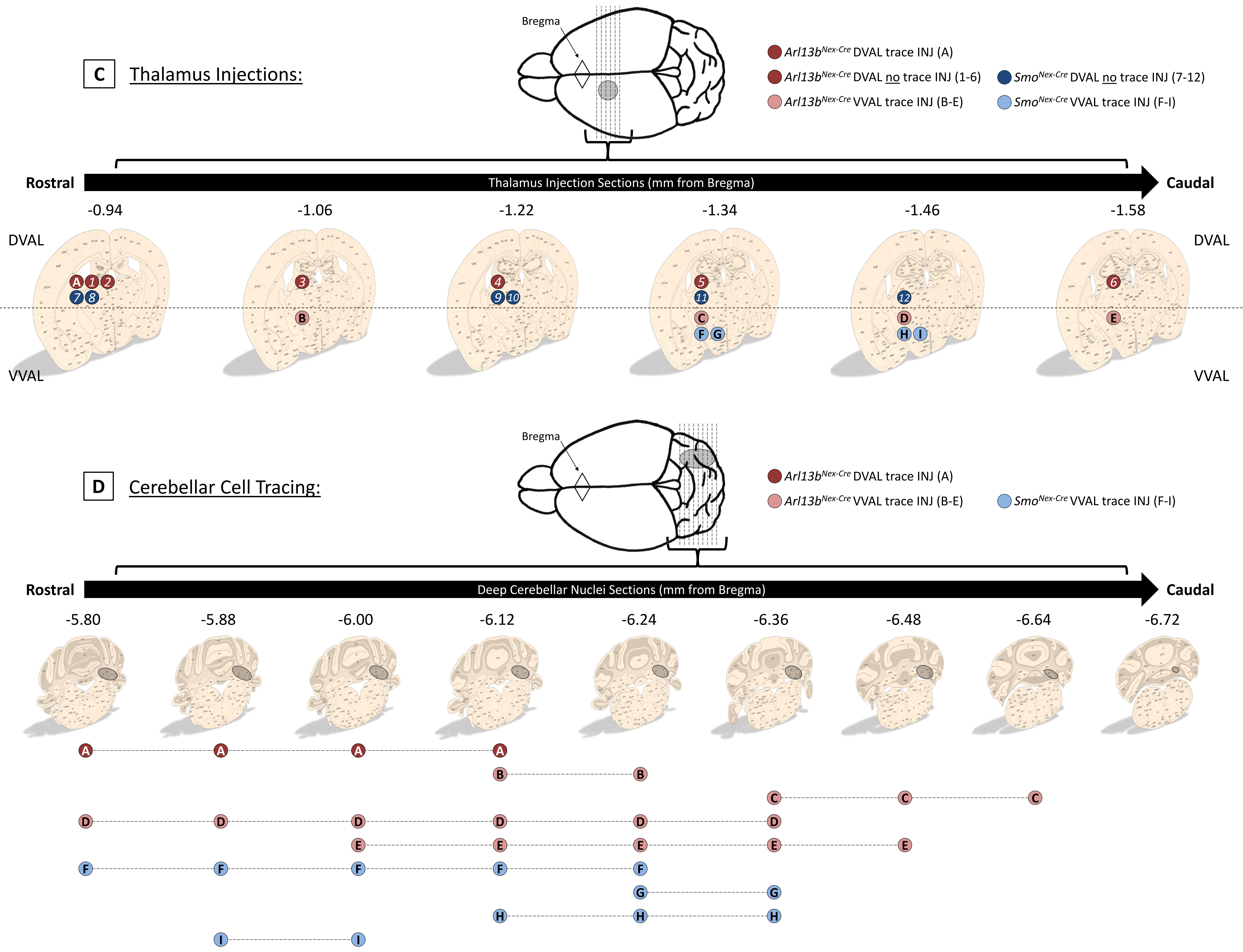
C
Thalamus Injections:
Bregma
Arl13b^Nex-Cre DVAL trace INJ (A)
Arl13b^Nex-Cre DVAL no trace INJ (1-6)
Arl13b^Nex-Cre VVAL trace INJ (B-E)
Smo^Nex-Cre DVAL no trace INJ (7-12)
Smo^Nex-Cre VVAL trace INJ (F-I)
Rostral
Thalamus Injection Sections (mm from Bregma)
Caudal
-0.94
-1.06
-1.22
-1.34
-1.46
-1.58
DVAL
VVAL
D
Cerebellar Cell Tracing:
Bregma
Arl13b^Nex-Cre DVAL trace INJ (A)
Arl13b^Nex-Cre VVAL trace INJ (B-E)
Smo^Nex-Cre VVAL trace INJ (F-I)
Rostral
Deep Cerebellar Nuclei Sections (mm from Bregma)
Caudal
-5.80
-5.88
-6.00
-6.12
-6.24
-6.36
-6.48
-6.64
-6.72

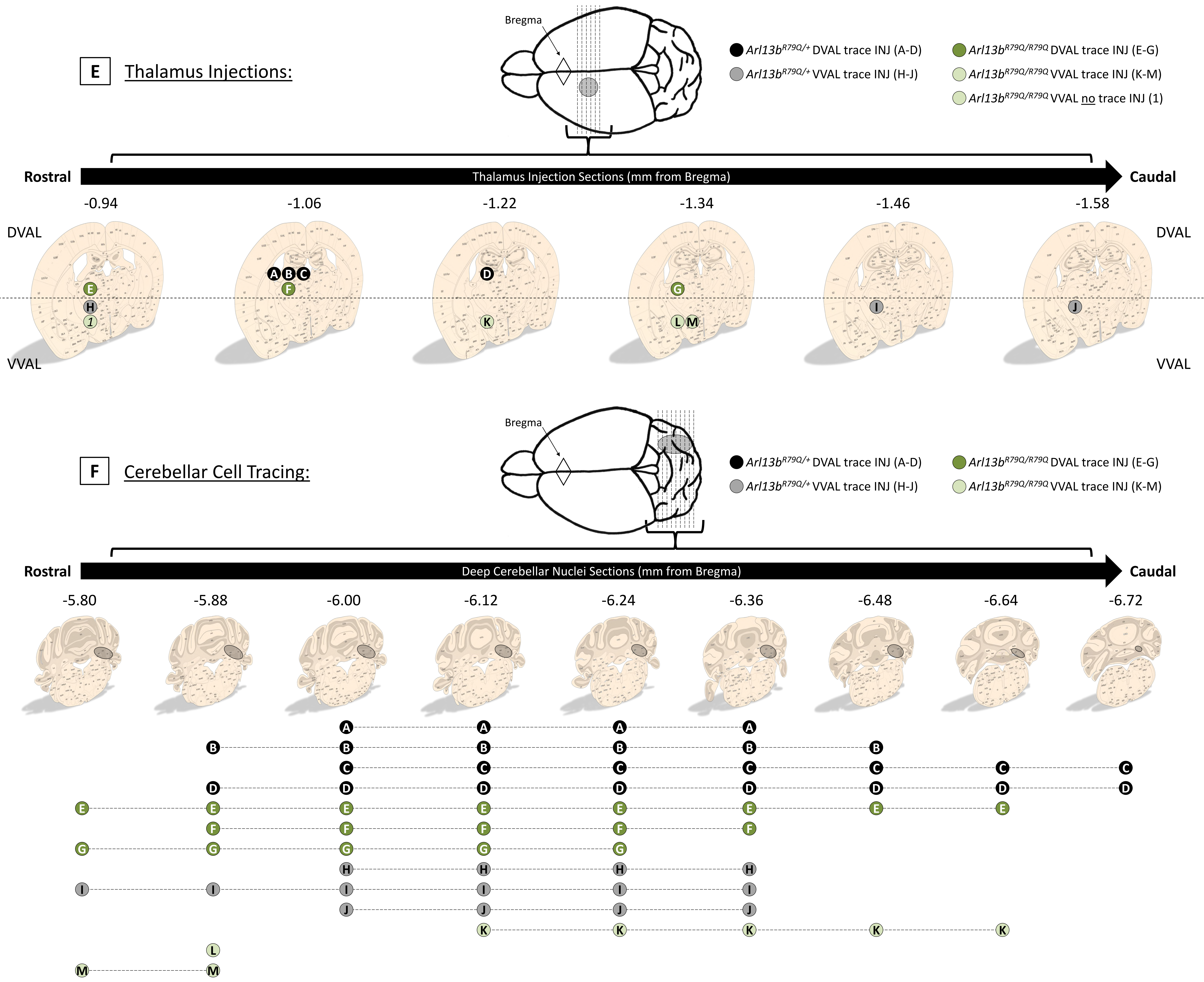
E
Thalamus Injections:
Bregma
*Arl13b*$^{R79Q/+}$ DVAL trace INJ (A-D)
*Arl13b*$^{R79Q/+}$ VVAL trace INJ (H-J)
*Arl13b*$^{R79Q/R79Q}$ DVAL trace INJ (E-G)
*Arl13b*$^{R79Q/R79Q}$ VVAL trace INJ (K-M)
*Arl13b*$^{R79Q/R79Q}$ VVAL no trace INJ (1)
Rostral
Thalamus Injection Sections (mm from Bregma)
Caudal
-0.94
-1.06
-1.22
-1.34
-1.46
-1.58
DVAL
VVAL
DVAL
VVAL
F
Cerebellar Cell Tracing:
Bregma
*Arl13b*$^{R79Q/+}$ DVAL trace INJ (A-D)
*Arl13b*$^{R79Q/+}$ VVAL trace INJ (H-J)
*Arl13b*$^{R79Q/R79Q}$ DVAL trace INJ (E-G)
*Arl13b*$^{R79Q/R79Q}$ VVAL trace INJ (K-M)
Rostral
Deep Cerebellar Nuclei Sections (mm from Bregma)
Caudal
-5.80
-5.88
-6.00
-6.12
-6.24
-6.36
-6.48
-6.64
-6.72

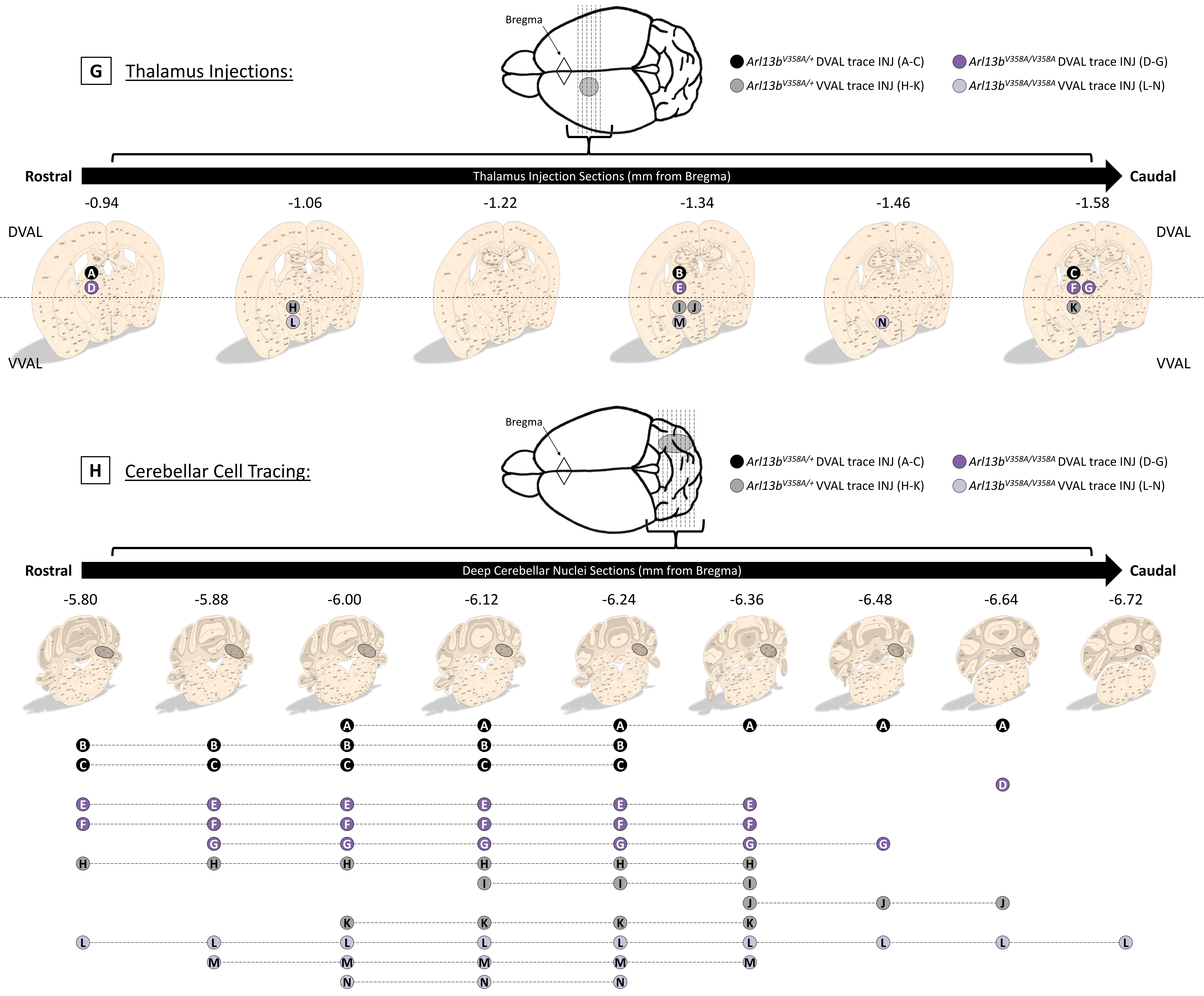
G
Thalamus Injections:
Bregma
Arl13b^V358A/+ DVAL trace INJ (A-C)
Arl13b^V358A/V358A DVAL trace INJ (D-G)
Arl13b^V358A/+ VVAL trace INJ (H-K)
Arl13b^V358A/V358A VVAL trace INJ (L-N)
Rostral
Thalamus Injection Sections (mm from Bregma)
Caudal
-0.94
-1.06
-1.22
-1.34
-1.46
-1.58
DVAL
VVAL
DVAL
VVAL
A D
H L
B E I J M
N
C F G K
H
Cerebellar Cell Tracing:
Bregma
Arl13b^V358A/+ DVAL trace INJ (A-C)
Arl13b^V358A/V358A DVAL trace INJ (D-G)
Arl13b^V358A/+ VVAL trace INJ (H-K)
Arl13b^V358A/V358A VVAL trace INJ (L-N)
Rostral
Deep Cerebellar Nuclei Sections (mm from Bregma)
Caudal
-5.80
-5.88
-6.00
-6.12
-6.24
-6.36
-6.48
-6.64
-6.72
A A A A A A
B B B B B
C C C C C
D
E E E E E E
F F F F F F
G G G G G G
H H H H H H
I I I
J J J
K K K K
L L L L L L L L L
M M M M M
N N N

**Figure S1: Total injection results.** (A, C, E, G) Thalamus injection site range. Shown at top are coronal sections of the cortex at the location of the thalamus, labeled with distance from Bregma (diamond) in mm. Circles represent injection sites as identified by needle mark, presence of dye, and tissue landmarks. Injections that resulted in fluorescent labeling of deep cerebellar nuclei (DCN) are denoted by letter, whereas injections that did not result in fluorescent DCN are denoted by number. Injections into the dorsal thalamus (DVAL) are represented as dark circles with white text; injections into the ventral thalamus (VVAL) are represented as light circles with black text. (B, D, F, H) Cerebellar cell tracing range. Shown at top are coronal sections where fluorescent DCN cell clusters were observed, labeled with distance from Bregma (diamond) in mm. Shown below are lines corresponding to the lettered injection sites (from the top half of each diagram) – the lines stretch to cover and mark the cerebellar sections in which fluorescent DCN were observed. Injection and tracing results for (A-B) control, (C-D) $Arl13b^{Nex\text{-}Cre}$ or $Smo^{Nex\text{-}Cre}$ mutant, (E-F) $Arl13b^{R79Q}$ control or mutant, and (G-H) $Arl13b^{V358A}$ control or mutant mice.
