## Supplementary material for "Smoothened and ARL13B are critical in mouse for superior cerebellar peduncle targeting": Supplmental figure 2

**A**

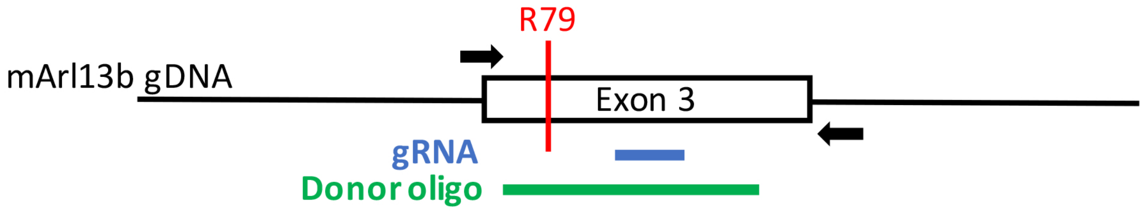


**B**

```
                   78   79   80   81   82   83   84   85   86   87   88   89   90   91
Wildtype      5'…ATT  CGG  GGA  ATT  TGG  AAG  AAT  TAT  TAT  GCT  GAA  TCC  TAT  GGG…3'
                  …I    R    G    I    W    K    N    Y    Y    A    E    S    Y    G…

Donor oligo   5'…ATT  CAG  GGC  ATA  TGG  AAG  AAT  TAT  TAT  GCT  GAA  TCC  TAT  GGG…3'
                        Q      NdeI  RE

CRISPR gRNA                                  5'  AT  TAT  TAT  GCT  GAA  TCC  TAT  GG  3'
                                                                             PAM
```

**Figure S2: *Arl13b*$^{R79Q}$ CRISPR design.** (A) Diagram showing the mouse Arl13b genomic DNA surrounding exon 3, with locations of the primers used for genotyping and sequencing (black arrows), guide RNA (gRNA, blue) and donor oligo (green) indicated. (B) Schematic of the 22-nucleotide CRISPR gRNA with the PAM sequence underlined, aligned with the *Arl13b* target sequence (wildtype). The 180bp donor oligo, containing regions of homology flanking each side of the target site, was used as a template for homology directed repair. This oligo contains the R-to-Q mutation (red) and two silent base changes (green) to add a *NdeI* restriction site (italicized).
